# Measuring Historical and Compositional Signals in Phylogenetic Data

**DOI:** 10.1101/2020.01.03.894097

**Authors:** Lars S Jermiin, Bernhard Misof

## Abstract

Most commonly-used molecular phylogenetic methods assume that the sequences evolved on a single bifurcating tree and that the evolutionary processes operating at the variable sites are Markovian. Typically, it is also assumed that these evolutionary processes were stationary, reversible and homogenous across the edges of the tree and that the multiple substitutions at variable sites occurred so infrequently that the historical signal (i.e., the signal in DNA that is due to the order and time of divergence event) in phylogenetic data has been retained, allowing for accurate phylogenetic estimates to be obtained from the data. Here, we present two metrics, *λ* and *δ*_*CFS*_, to quantify the strength of the historical and compositional signals in phylogenetic data. *λ* quantifies *loss of historical signal*, with *λ* = 0.0 indicating evidence of a strong historical signal and *λ* = 1.0 indicating evidence of a fully eroded historical signal. *δ*_*CFS*_ quantifies *compositional distance* from full symmetry of a divergence matrix generated by comparing two sequences, with *δ*_*CFS*_ = 0.0 indicating no evidence of evolution under dissimilar conditions and *δ*_*CFS*_ > 0.0 indicating increasing evidence of lineages diverging under different conditions. The metrics are implemented in methods intended for use after multiple sequence alignment and before model selection and phylogenetic analysis. Results generated using these methods allow users of phylogenetic tools to select phylogenetic data more wisely than it previously was possible. The merits of these metrics and methods are illustrated using simulated data and multi-gene alignments obtained from 144 insect genomes.

---

Model-based molecular phylogenetic methods typically assume that the sequences of nucleotides or amino acids have diverged over the edges of a bifurcating tree, and that the evolutionary processes that operate at the variable sites of these data (i.e., sites that are free to change) can be approximated by independent, identically-distributed Markov processes. Also, it is assumed that the evolutionary processes were stationary, reversible, and homogeneous (SRH)—for details, consult Bryant et al. (2005), Jayaswal et al. (2005), Ababneh et al. (2006a,b) and Jermiin et al. (2017, 2020a). These assumptions apply equally to single-gene, concatenated-gene, and gene tree/species tree approaches in phylogenetics. The only difference is that the topologies of gene trees are assumed to be identical in the concatenated-gene approach but are free to vary in the gene tree/species tree approach.

Inferring phylogenetic estimates from sequence data is aided by a growing number of phylogenetic programs, including PAUP* (Swofford 2002), PHYLIP (Felsenstein 2005), PhyML (Guindon et al. 2010), MrBayes (Ronquist et al. 2012), PhyloBayes (Lartillot et al. 2013), Garli (Bazinet et al. 2014), RAxML (Stamatakis 2014), HAL-HAS (Jayaswal et al. 2014), IQ-TREE (Nguyen et al. 2015) and BEAST (Bouckaert et al. 2019). All of these methods accommodate the fact that multiple substitutions at the same variable site erase all evidence of previous substitutions at these sites. However, there is a limit to how far apart sequences can diverge before the *historical signal* (i.e., the signal in DNA that is due to the order and timing of divergence events) has eroded so much that it is pointless attempting to infer a phylogeny from these data.

Visually, an alignment may appear conserved, but that is because we tend to focus on the presence and abundance of constant sites in the alignment. If we were to focus our attention on the differences at the variable sites, we may actually find that the nucleotides at these sites have changed so frequently that the historical signal is erased. Unfortunately, while it is well known that multiple substitutions at the same sites will erode a historical signal in an alignment (Ho and Jermiin 2004) and, therefore, that there is a *phylogenetic twilight zone* (Chang et al. 2008), little has been done to quantify how eroded a historical signal in an alignment of sequence data may be. An exception to this is Xia et al.’s (2003) index of substitution saturation. Using an entropy-based approach, this method computes the average entropy per site, 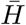, for the alignment and compares it to the expected value, *H*_*FSS*_, which is computed by assuming that the nucleotide frequencies at each site can be drawn from a multinomial distribution. Computer simulations suggest that the index is a reliable estimator of the decay of a phylogenetic signal (Xia et al. 2003), but a closer look at the underpinning algebra shows that it is not suitable for sequences that have diverged under complex evolutionary conditions (e.g., non-homogeneous conditions). Furthermore, it assumes that all sites are variable, so the method may yield misleading results, even if the estimates of 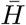 and *H*_*FSS*_ are derived from just the variant sites.

Here, we propose a method that quantifies the *loss of historical signal* from pairs of sequences that may have acquired different nucleotide frequencies since they diverged from their last common ancestor, and the presence of invariable sites is allowed for. In addition, we explore a recently proposed *standardised compositional distance* (Jermiin et al. 2020a) as a means of measuring *emergence of compositional signals* from pairs of sequences. Here, the compositional signal is one of the *non-historical signals* that might arise over time due to lineage-specific differences in the evolutionary process (Jermiin et al. 2020a).

Within the phylogenetic protocol (Jermiin et al. 2020b), the methods are intended for use after the initial multiple sequence alignment has been completed and before model selection is commenced. The methods are accurate, fast to use, and easy to employ.

## MATERIALS AND METHODS

### Three Useful Metrics

Consider a multiple sequence alignment (**A**) of nucleotides with *a* sequences and *b* variable sites. This set of sites includes all the variant sites (i.e., those displaying evidence of having changed). For each sequence pair [*i, j*], obtain a 4 × 4 divergence matrix, **N**:

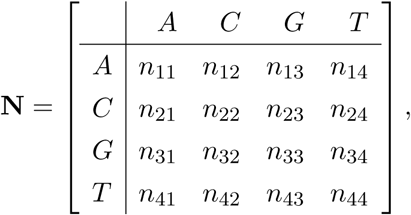

where *n*_*kl*_ represents the number of sites with nucleotide *k* in sequence *i* and nucleotide *l* in sequence *j* (here, the indices *k, l* = 1, …, 4, correspond to the nucleotides *A, C, G*, and *T*). Furthermore, let *n*_*k*•_ =∑_*l*_ *n*_*kl*_, *n*_•*l*_ =∑_*k*_ *n*_*kl*_ and *n*_••_ =∑_*k,l*_ *n*_*kl*_. Finally, let

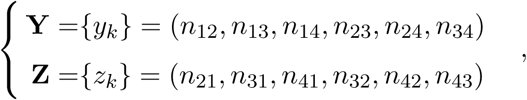

where *m* = 1, …, 6 (here, *m* is the number of elements in **Y** and **Z**).

Given these definitions, we can compute:

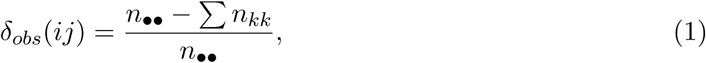

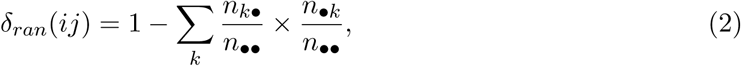

and

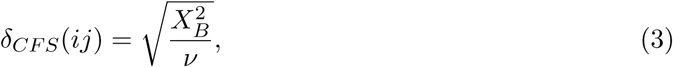

where *δ*_*obs*_(*ij*) is the proportion of variable sites with different nucleotides in sequences *i* and *j, δ*_*ran*_(*ij*) is the proportion of variable sites with different nucleotides in sequences *i* and *j*, assuming they have evolved for infinitely long time under independent stationary conditions—which implies that each sequence will be in equilibrium—and *δ*_*CFS*_ (*ij*) is a *standardised compositional distance* between two vectors containing compositional data (i.e., **X** and **Y**). Equation (4) is obtained from the matched-pairs test of symmetry by Bowker (1948), where 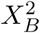 is the test statistic and *ν* is the number of degrees of freedom (Jermiin et al. 2020a). Given these metrics, three distance matrices (i.e., **D**_*obs*_, **D**_*ran*_ and **D**_*CFS*_) can be computed from **A**, allowing us to survey the historical and compositional signals from pairs of sequences in phylogenetic data.

### Measuring the Decay of a Historical Signal

To determine whether the historical signal between two sequences has been lost, we define the *loss of historical signal* between two sequences as:

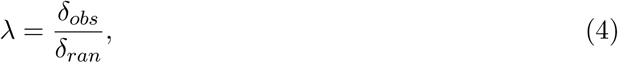

where *λ* ≥ 0.0. If the nucleotides at the variable sites are identical, *λ* = 0.0 and there is no evidence of a loss of historical signal between the sequences. On the other hand, if the two sequences are random with respect to each other, *λ* = 1.0 and the historical signal will be fully decayed. Occasionally, *λ* > 1.0, in which case *δ*_*obs*_ or *δ*_*ran*_ may be inaccurate, perhaps due to a finite length of the alignment or due to sequencing or alignment error.

The main advantage of this method of estimating loss of historical signal is that *λ* allows for cases where the sequences have diverged under different Markovian conditions; hence, *λ* is less likely to be overestimated. Likewise, *δ*_*obs*_ and *δ*_*ran*_ can be obtained from all sites in an alignment or just the variant sites in the alignment. This allows the investigator to decide whether the constant sites should be regarded as variable or invariable. That decision has to be made before any survey is begun and whatever is decided will apply to all of the 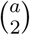 sub-alignments.

### Measuring the Emergence of a Compositional Signal

It is well known that the historical signal in a sequence alignment decays over time and that the strength of the non-historical signals (e.g., due to compositional heterogeneity arising across the sequences) may increase over time (Ho and Jermiin 2004). It seems less widely known that the historical and non-historical signals might interact synergistically (i.e., the signals align, supporting each other) and antagonistically (i.e., the non-historical signals may conceal the historical signal, making it difficult to extract the historical signal using commonly-used phylogenetic methods) in a phylogenetic analysis (Ho and Jermiin 2004).

Obviously, inspecting the inferred phylogenetic tree does not allow us to conclude whether the estimate is biased by non-historical signals. However, it is possible to assess whether there is an association between the values of *λ* and *δ*_*CFS*_. If such an association exists (e.g., with high values of *λ* concurring with high values of *δ*_*CFS*_), then there might be cause for concern because it means that lineage-specific differences in the evolutionary process may have arisen. These differences might bias estimates of phylogenetic trees (Ho and Jermiin 2004; Jermiin et al. 2004) unless they are modelled optimally in terms of the parameters assumed to be necessary.

### New Tools for Surveying Phylogenetic Data

To enable the surveys described above, three newly-released programs are available from:

- http://www.github.com/lsjermiin/SatuRation.v1.0/
- http://www.github.com/ZFMK/SatuRationHeatMapper/
- http://www.github.com/ZFMK/RedundancyHeatMapper/

and are described below.

#### SatuRation

SatuRation v1.0 is a data-surveying program written in C++ and is released under a GNU General Public License v3.0. It computes *δ*_*obs*_, *δ*_*CFS*_ and *λ* for all pairs of sequences in an alignment and can be executed using the following commands:

~~~
saturation <infile> <a|v> <b|f> <1|…|31>
~~~

or

~~~
saturation <infile> <a|v> <b|f> <1|…|31> > README
~~~

where `infile` is a text file with an alignment of characters in the fasta format, `a|v` refers to whether all sites or just the variant sites should be used, `b|f` refers to whether a brief or full report of the results should be printed, and `1|…|31` refers the data type and how the data should be analysed. If `b` is used, SatuRation prints a line with key statistics to the user’s terminal; if `f` is used, it prints six files with the values of *δ*_*obs*_, *δ*_*CFS*_, and *λ*, three files in the `.dis` format and three files in the `.csv` format. Likewise, if `v` is used, it prints a file with an alignment of the variant sites used. A summary of the results is also printed to the terminal or to the `README` file (doing the latter is useful if multiple data sets are surveyed).

SatuRation is designed to analyse alignments of nucleotides, di-nucleotides, codons, 10- and 14-state genotypes, and amino acids. If the `infile` contains sequences of:

- Single nucleotides (4-state alphabet), the sequences may be recoded into six 3-state alphabets or seven 2-state alphabets,
- Di-nucleotides (16-state alphabet; i.e., *AA, AC*, …, *TG, TT*), the sequences may be divided into alignments with 1st or 2nd position sequences,
- Codons (a 64-state alphabet; i.e., *AAA, AAC*, …, *TTG, TTT*), the sequences may be divided into three alignments with di-nucleotide sequences and three alignments with single-nucleotide sequences,
- Amino acids (a 20-state alphabet), the letters may be recoded to a 6-state alphabet. This type of recoding was recently used to study early evolution of animals (Feuda et al. 2017). Other types of recoding amino acids have been used (Kosiol et al. 2004; Susko and Roger 2007) but are not considered.

The 10- and 14-state genotype data cater for diploid and triploid genomes. For example, if a locus in a diploid genome contains nucleotides *A* and *G*, then the genotype sequence will contain an *R* at that locus. There are 10 distinguishable genotypes for each locus in diploid genomes and 14 for every locus in triploid genomes. For further detail about the data types and how the data may be analysed, simply type:

~~~
saturation
~~~

on the command line and follow the instructions.

The output from SatuRation falls into two categories: `.csv` files and `.dis` files. The `_table.csv` file contains the estimates obtained for all pairs of sequences. It can be opened and viewed by, for example, Microsoft Excel. The `_dobs.csv`, `_dcfs.csv` and `_lambda.csv` files, respectively, contain the *δ*_*obs*_, *δ*_*CFS*_, and *λ* values, set out in a format that can be read by SatuRationHeatMapper and RedundancyHeatMapper (see below). The three `.dis` files contain the *δ*_*obs*_, *δ*_*CFS*_, and *λ* values, and can be analysed further using TreeLikeness (see below), FastME (Lefort et al. 2015) and SplitsTree4 (Bryant and Moulton 2004). Finally, the `_used.fst` file contains an alignment of the sites used; this alignment may be necessary in other analyses not considered here.

#### SatuRationHeatMapper

SatuRationHeatMapper v1.0 is designed to generate a color-coded heat map from the `_lambda` file. The colors used range from white (*λ* < 0.64) to black (*λ* ≤ 1.0), with *λ* > 1.0 being highlighted using the color red. SatuRationHeatMapper is written in Perl and can be executed using the following command:

~~~
SatuRationHeatMapper -i <infile> -<t|f>
~~~

where `infile` must be the `_lambda.csv` file and where `t` and `f` stand for triangle and full, respectively. The output is an `.svg` file with a heat map in scalable vector graphics format. This file can be opened and processed using Adobe Illustrator.

#### RedundancyHeatMapper

RedundancyHeatMapper v1.0 is designed to generate a color-coded heat map from the `_lambda.csv` file. The colors used range from white (*λ* < 0.01) to black (*λ* ≥ 0.29). RedundancyHeatMapper is written in Perl and can be executed using the following command:

~~~
RedundancyHeatMapper -i <infile> -<t|f>
~~~

where `infile` must be the `_lambda.csv` file and where `t` and `f` stand for triangle and full, respectively. The output is an `.svg` file with a heat map in scalable vector graphics format. This file can be opened and processed using Adobe Illustrator.

## RESULTS

### Measuring Decay of the Historical Signal

To illustrate the merits of *λ*, we examined an alignment of 12 nucleotide sequences generated by simulation under the F81 model of nucleotide substitutions (Felsenstein 1981) on the tree outlined in Figure 1a. All sites had the same probability of change. In practice, the data were generated using INDELible (Fletcher and Yang 2009). Given this tree (Fig. 1a), we expected the distribution of *λ* values to be penta-modal (i.e., with one peak based on comparisons between `SeqI`, `SeqJ`, `SeqK`, and `SeqL`, another peak based on comparisons between `SeqE`, `SeqF`, `SeqG`, and `SeqH`, and so forth).

**Figure 1:**
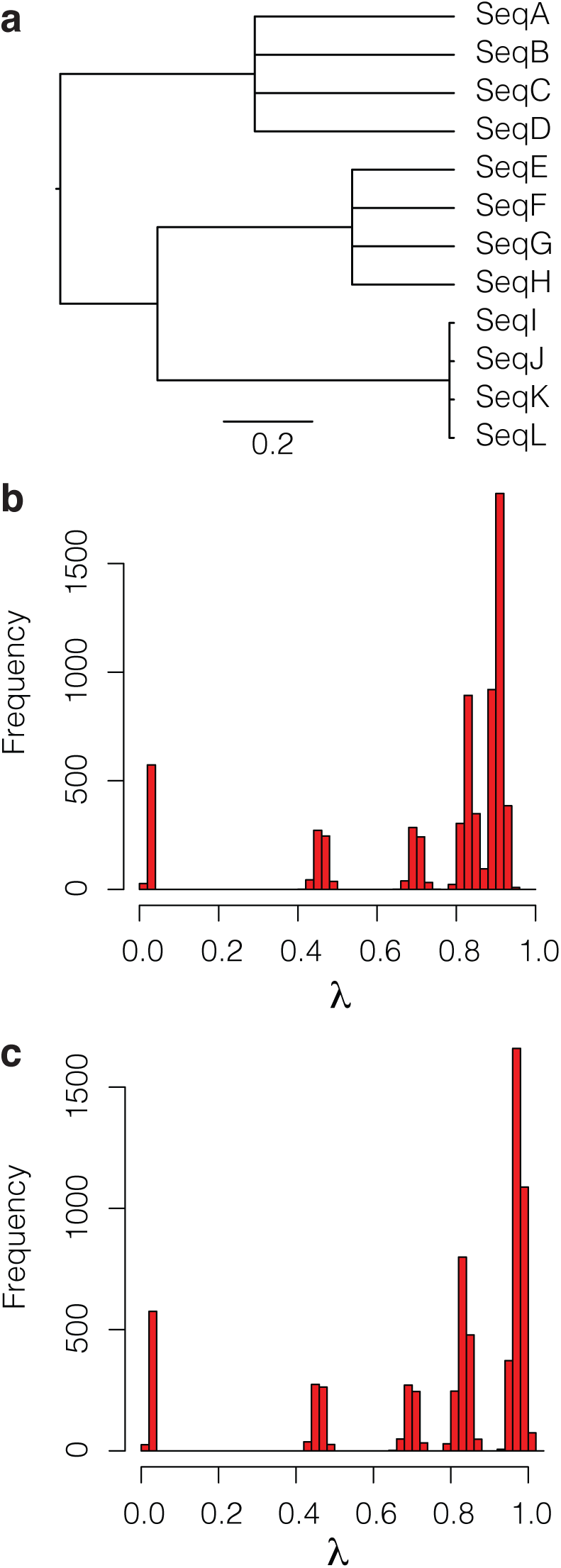
Diagram showing (**a**) the 12-tipped tree used to simulate alignments of nucleotides with 2500 sites, and two histograms (**b** and **c**) with the distributions of *λ* values (based on 100 replicates). The bar in **a** corresponds to 0.2 substitution per site. In **b**, the root-to-tip distance was 0.89 while in **c**, it was 1.39. All other divergences were kept the same. Sequences were generated using INDELible (Fletcher and Yang 2009).

The histogram in Figure 1b, obtained by pooling the *λ* values from 100 data sets, reveals the expected penta-modal distritution. The left-most peak is due to comparisons between `SeqA`, `SeqB`, `SeqC`, and `SeqD`, the next peak is due to comparisons between `SeqE`, `SeqF`, `SeqG`, and `SeqH`, and so forth. As expected, although the five divergence events are separated by the same interval of time (i.e., they occurred 0.89, 0.67, 0.45, 0.23, and 0.01 substitutions ago), the distance between the neighboring peaks is largest for low values of *λ* and smallest for high values of *λ*. Interestingly, none of the 6,600 estimates in Figure 1b reached 1.0, implying that none of the sequences can be regarded as random with respect to each other. In other words, while the historical signal is quite eroded for many pairs of sequences (notably, comparisons where `SeqA`, `SeqB`, `SeqC` or `SeqD` is one of the sequences), it is not yet completely decayed.

A second experiment was done to assess the effect of moving the initial divergence back in time from 0.89 to 1.39. The result of doing so was insightful. The righthand peak moved further to the right whereas the other peaks remained in the same locations (Fig. 1c). The righthand peak now covers *λ* = 1.0, implying a complete decay of the historical signal for some of the sequences (i.e., comparisons where `SeqA`, `SeqB`, `SeqC` or `SeqD` is one of the sequences). In other words, displaying the values of *λ* obtained from alignments of nucleotides may be a useful first step in assessing usefulness of phylogenetic data.

### Informed Removal of Sequences

Having found that there might be a problem (i.e., a decayed historical signal) with some of the sequences in a phylogenetic data set, there is the issue of identifying the *most affected sequences*—if they can be named, then the intended phylogenetic analysis may be done with or without them.

The heat map in Figure 2a shows the colour-coded *λ* values for each sequence pair. In this example, the focus is on *λ* values above 0.64, so the map is called a *saturation plot*. The saturation plot shows that the highest values of *λ* are between one group, consisting of `SeqA`, `SeqB`, `SeqC`, and `SeqD`, and another group, comprising the remaining eight sequences. Obviously, a saturation plot may be used to identify the sequences most likely to have lost their historical signal (with respect to each other).

**Figure 2:**
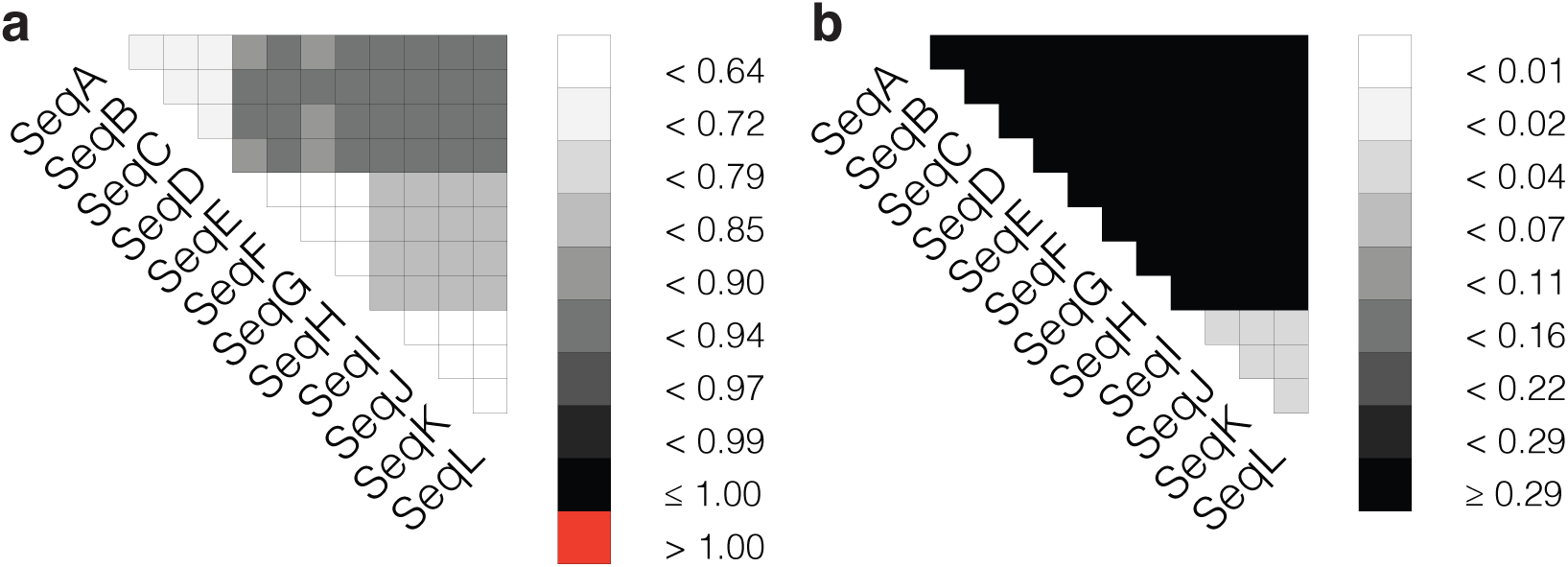
Saturation plot (**a**) and redundancy plot (**b**). The two heat maps were obtained for one of the 100 replicates referred to in Fig. 1.

Sometimes an alignment may contain more sequences than we are willing (or able) to handle. This situation may arise if a gene tree is to be inferred by maximum-likelihood or Bayesian methods. Then, it is important to bear in mind that every sequence added to the alignment entails having to estimate at least two more parameters (the edge lengths), and that it is recommended that sample size exceeds 40 times the number of parameters that must be optimised (Burnham and Anderson 2002). In molecular phylogenetics, this number (i.e., sample size) is determined by the number of sites in an alignment (Posada and Crandall 2001), which is typically less than a few thousand sites for alignments used to infer gene trees. The heat map in Figure 2b shows the colour-coded value of *λ* for all pairs of sequences. In this case, the focus is on *λ* values below 0.29, so the map is called a *redundancy plot*. The redundancy plot reveals sequences that are so similar to one another that removal of some of the sequences may be worth considering. Doing so is unlikely to affect the phylogenetic estimate, but it will lower the number of parameters that must be optimised and that can lead to smaller variances and faster phylogenetic analyses.

Prior to generating a saturation plot, it is recommended that the constant sites be removed from the alignment. This is because it is impossible to say whether constant sites are invariable (i.e., cannot change) or simply have not changed. Removal of constant sites may inflate the estimates of *λ*, but since it is the variant sites that determine the optimal shape of a phylogenetic tree, being alerted to cases of a highly-decayed historical signal is preferable. On the other hand, if the objective is to reduce the number of sequences in an alignment, then the constant sites can be retained in the alignment.

### Measuring the Emergence of a Compositional Signal

Figure 3 shows a dot plot, with the *δ*_*CFS*_ value as a function of the *λ* value, for all pairs of sequences surveyed in Figure 1. The penta-modal distributions of *λ* in Figures 1b and 1c are replicated in Figure 3. An interesting, but also unexpected, feature of Figure 3 is that the five distributions of *δ*_*CFS*_ appear to be largely independent of the value of *λ*.

**Figure 3:**
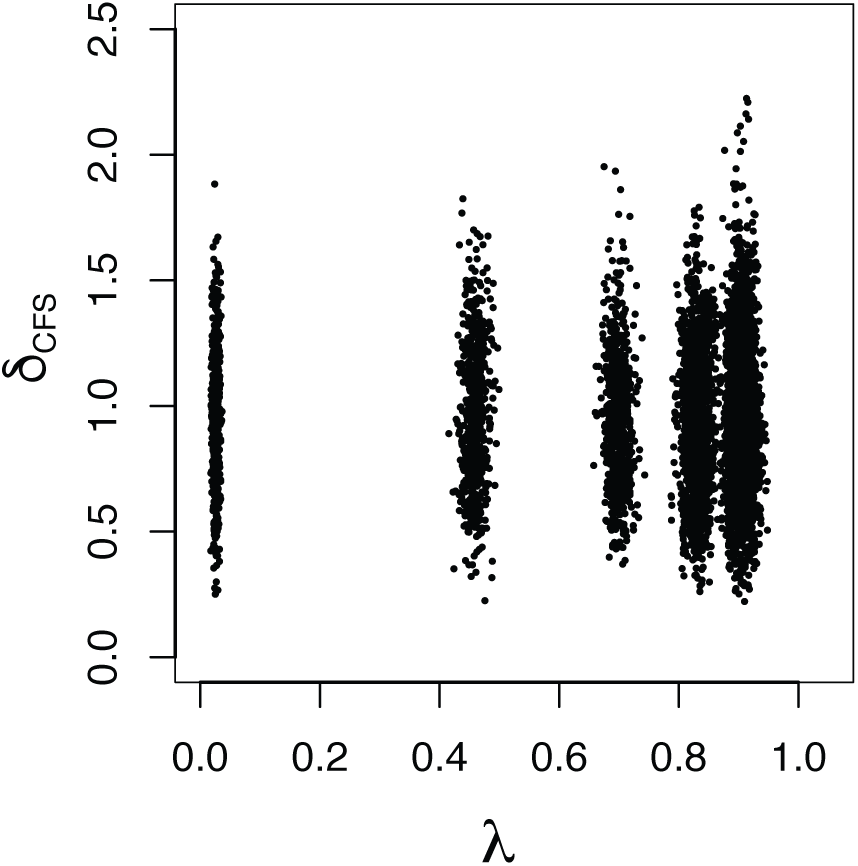
Dot plot showing the *δ*_*CFS*_ values as a function of the *λ* values for the sequences analysed in Figure 1.

To determine whether this feature represents what to expect under a wider range of tree topologies, we repeated the experiment using 25-tipped, random trees simulated under a birth-death process (Yang and Runnala 1997). In practice, the trees were obtained using evolver, from the PAML program package (Yang 2007), with the birth rate, death rate and sampling fraction drawn at random from a continuous uniform distribution on the interval [0,1), and with the tree depth set to 0.5, 1.0, 2.0, or 4.0. Twenty-five trees were generated for each tree depth. For each tree, we simulated an alignment of 2500 nucleotides under the F81 model. Data were generated using INDELible (Fletcher and Yang 2009). From each of these alignments, we obtained 300 estimates of *δ*_*CFS*_, one for each sequence pair.

Table 1 presents the mean, variance, minimum and maximum of *δ*_*CFS*_ derived from the alignment generated in this experiment. There is no discernible difference among these four distributions of *δ*_*CFS*_ values, except that there might be a subtle rise in the maximum value of *δ*_*CFS*_ as the tree depth increases from 0.5 to 4.0. However, this increase is unlikely to have any practical consequences, because the historical signal has already decayed fully (*λ* = 1.0) for 22% and 45% of the sequence pairs obtained on trees with tree depths of 2.0 and 4.0, respectively (0% was obtained for trees with tree depths of 0.5 and 1.0).

**Table 1:** Summary statistics for *δ*_*CFS*_ based on computer simulations on 25-tipped trees with different tree depth. For each tree depth, the mean, variance, minimum, and maximum of *δ*_*CFS*_ is shown, along with the number of sequence pairs being compared.

| Tree depth | Mean | Variance | Min. | Max. | Pairs |
| --- | --- | --- | --- | --- | --- |
| 0.5 | 0.9525 | 0.0760 | 0.1433 | 1.9716 | 7500 |
| 1.0 | 0.9474 | 0.0813 | 0.1100 | 2.0728 | 7500 |
| 2.0 | 0.9683 | 0.0827 | 0.1997 | 2.2274 | 7500 |
| 4.0 | 0.9640 | 0.0796 | 0.1929 | 2.2410 | 7500 |

This result has two implications. If a survey of real data returns: (a) a distribution of *δ*_*CFS*_ that differs markedly from those in Table 1, then there may be evidence that some of the sequences have evolved under different conditions, and (b) a maximum value of *δ*_*CFS*_ greater than 2, then the sequences in the corresponding sequence pair may be among those most likely to have evolved under different conditions.

### Example: A Pre-phylogenetic Survey of Phylogenetic Data from Insects

The merits of the methods presented above are highlighted in Figure 4. It shows the values of *δ*_*CFS*_ as a function of the *λ* values for the nucleotide sequences that underpin the phylogenetic estimate of the evolution of 144 species of insects (Misof et al. 2014). The dot plots, one for each codon site, differ markedly from one another but concur in two regards. For these data: (a) the historical signal is never fully decayed for any pair of sequence, even when the data comprise third codon sites (*λ*_*max*_ = 0.9459), and (b) the summary statistics for *δ*_*CFS*_ differ vastly from those in Table 1, with most values being greater than 2. Hence, for these data, a preliminary survey of the phylogenomic data would lead to the following conclusions:

**Figure 4:**
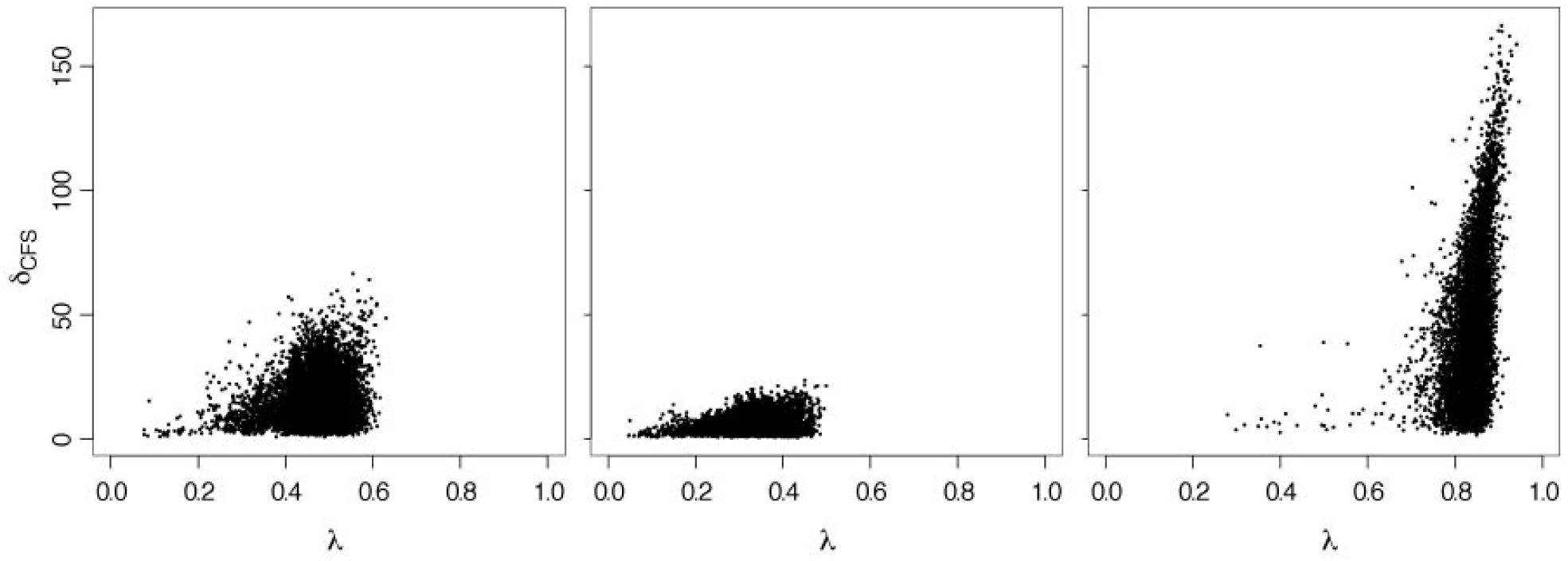
Dot plots showing the *δ*_*CFS*_ values as a function of the *λ* values for first (left), second (centre), and third (right) codon sites from an alignment with 144 sequences and 413,459 codons. The alignments were first used by Misof et al. (2014). Only variant sites were considered. The summary statistics (mean, variance, minimum, and maximum) for *δ*_*CFS*_ are: first codon sites (14.8409, 87.7655, 0.7649, 66.5152); second codon sites (5.2480, 9.5935, 0.5160, 23.5955); third codon sites (38.6445, 746.3297, 1.3870, 166.2780).

- None of the sequences in alignments of 1st, 2nd or 3rd codon sites can be regarded as random with respect to each other, so the historical signal is clearly not eroded fully, even though the signal is very low, as expected, at 3rd codon sites.
- Most of the sequences in the alignments of 1st, 2nd or 3rd codon sites appear to have evolved under different Markovian conditions, so a more rigorous assessment of these alignments with the matched-pairs test of symmetry (Ababneh et al. 2006b; Jermiin et al. 2020a) is required. Indeed, such an assessment was done by Misof et al. (2014), who discovered that most of the sequences in their alignments of genes probably had evolved under non-homogeneous conditions.

## DISCUSSION

In phylogenetics, the historical signal in sequence data is known to decay over time (Ho and Jermiin 2004). Conversely, the compositional signal in these data might increase over time. That both of these signals might be present in phylogenetic data leaves anyone interested in inferring accurate phylogenetic estimates from the data with a big challenge. They can either proceed—as most published phylogenetic studies have done so far—and assume that violation of the phylogenetic assumptions, while unavoidable, have no effect on the phylogenetic estimate, *or* they can employ a growing body of data-surveying tools intended for phylogenetic data, and gain as much information as possible about the data before they embark on the actual, sometimes very time-consuming, phylogenetic analysis.

The methods presented here are intended to fill a gap in our phylogenetic protocol (Jermiin et al. 2020b) between the multiple sequence alignment and the model selection. The gap exists because commonly-used phylogenetic methods are based on assumptions, and if they are violated by the data, there is an elevated risk of error in our phylogenetic estimates. Hence, it makes sense to survey phylogenetic data for evidence of violation of the phylogenetic assumptions *before* phylogenetic programs are applied to the data, but such pre-phylogenetic surveys are rarely done, and that is a worry because under certain conditions it is possible to infer the correct tree topology even though the historical signal is fully decayed (Ho and Jermiin 2004). Clearly, under such conditions the non-historical signals will dominate the historical signal, and any tree inferred from such data will be of little value to those hoping to infer the evolutionary history of species.

One of these assumptions concerns the historical signal in phylogenetic data. If the historical signal is heavily eroded, then other phylogenetic signals—like the compositional signal—may be strong enough to return a consistent phylogenetic estimate. However, this estimate might not reflect the true evolutionary history and there is no way of finding this out, unless parametric bootstrapping or predictive posterior probability analyses are done. These analyses are computationally expensive and time-consuming, so it is much better to examine the phylogenetic data for evidence of violation of phylogenetic assumptions before model selection than at a later stage in the phylogenetic protocol.

The method to estimate *λ* is a sensible solution to the problem of measuring decay of the historical signal. It relies on *δ*_*obs*_ (otherwise known as the *p* distance) and the metric *δ*_*ran*_, which corresponds to the proportion of variable sites with differences between the two sequences that have been allowed to evolve for infinitely long time under independent (and possibly different) Markovian conditions. These conditions may be such that the sequences will acquire different nucleotide compositions at the homologous sites. That *δ*_*ran*_ is flexible is desirable, because it is well known that homologous sequences often differ in nucleotide or amino-acid composition (Naser-Khdour et al. 2020). It is for this reason that *λ* may be more informative than the index proposed by Xia et al. (2003). Another reason is that *λ* applies to a pair of sequences whereas the index proposed by Xia et al. (2003) applies to the alignment of all the sequences. Using *λ*, it is possible to identify and delete sequences from an alignment on grounds that the historical signal is too eroded for comfort, or on grounds that a cluster of sequences are so similar to one another that it may be pointless to include them all in a phylogenetic analysis (e.g., Tay et al. (2017)). IQ-TREE (Nguyen et al. 2015) already removes all but two of the identical sequences before a phylogenetic analysis is commenced, but any sequence that differs from the other sequences at just one site will be kept in the alignment. The combined approach implemented in the methods described above facilitates naming all of the identical and near-identical sequences in an alignment; this means that anyone who uses phylogenetic methods to annotate genes will be able to work faster and obtain more accurate phylogenetic estimates during the gene annotation process.

The method to estimate *δ*_*CFS*_ is a sensible solution to the problem of estimating a standardised compositional distance between two sequences. Not only are all the constant sites in the data ignored, but the metric is also scaled such that estimates associated with different degrees of freedom can be compared directly. The benefit of this is illustrated in Figure 4 and by comparing the maximum values of *δ*_*CFS*_ from 1st (66.5152), 2nd (23.5955) and 3rd (166.2780) codon positions to that obtained from the corresponding alignments of amino acids (6.7964) and codons (10.6511). Because the size of the alphabet used equals *ν* in equation (4), we are now able to conclude that the maximum compositional distance is greatest for the alignment of 3rd codon sites and smallest for the alignment of amino acids. Had these results been available during the first analysis of these data (Misof et al. 2014), they could have been used to inform the strategy used to analyse the phylogenomic data phylogenetically. For example, more importance might have been put on the alignment of amino acids and less on the alignments of 1st and 2nd codon positions.

Finally, it came as a surprise that the historical signal is not completely eroded for 3rd codon position of most of the sequences (Fig. 4). This was unexpected because of the long time scale over which these sequences have evolved, and it is now clear that such an expectation might be ill-guided in many other instances where vertebrate or invertebrate nuclear DNA is compared phylogenetically. The matter that needs more attention is the analysis of compositionally heterogenous sequences because compositional signals in such data will be regarded as a historical signal by most currently-used phylogenetic methods.

## ACKNOWLEDGEMENTS

We wish to thank M. Hiller, G. Hughes and S.A. Shilling for constructive feedback.

